# Integration of spatially opposing cues by a single interneuron guides decision making in *C. elegans*

**DOI:** 10.1101/2023.01.23.525194

**Authors:** Asaf Gat, Vladyslava Pechuk, Sonu Peedikayil-Kurien, Gal Goldman, Jazz Lubliner, Shadi Karimi, Michael Krieg, Meital Oren-Suissa

## Abstract

The capacity of animals to integrate and respond to multiple hazardous stimuli in the surroundings is crucial for their survival. In mammals, complex evaluations of the environment require large numbers and different subtypes of neurons. The nematode *C. elegans* avoid hazardous chemicals they encounter by reversing their direction of movement. How does the worms’ compact nervous system processes the spatial information and directs the change of motion? We show here that a single interneuron, AVA, receives glutamatergic excitatory signals from head sensory neurons and glutamatergic inhibitory signals from the tail sensory neurons. AVA integrates the spatially distinct and opposing cues, whose output instructs the animal’s behavioral decision. We further find that the differential activation of AVA from the head and tail stems from distinct anatomical localization of inhibitory and excitatory glutamate-gated receptors along the AVA process, and from different threshold sensitivities of the sensory neurons to aversive stimuli. Our results thus uncover a cellular mechanism that mediates spatial computation of nociceptive cues for efficient decision-making in *C. elegans*.

## INTRODUCTION

Animals sense their surroundings by integrating multiple sensory cues and then translating them into motor actions. Smell, vision, touch, and proprioception all require that the animal detect, recognize, make a decision and respond to multiple sources of sensory information. The decision making is dependent on the prior processing of the perceived environmental cues^1–4^. For example, in the case of locomotion-related decisions, such as navigation, determining the spatial orientation is crucial for avoiding hazards or approaching a mating partner^5–7^. The computation of spatial sensorimotor information relies on the capacity of interneurons to integrate sensory inputs from multiple spatially distinct sources^1,5^. This type of integration can enhance the salience of stimuli and hasten behavioral responses^1^. Conceptually, spatial decision-making should be governed by a mechanism that can perform the necessary complex comparison and computation of spatially distinct inputs in order to determine the optimal behavioral output^8^. Previous studies confirmed that the integration of spatial information in mammals is encoded at the neural population level within specific brain structures, such as the superior colliculus^9,10^. However, can a single neuron integrate and execute spatial decision making? What are the molecular pathways, synaptic properties and neuronal activity patterns that mediate this process?

*C. elegans* exhibits a diverse repertoire of orientation behaviors to locate food, mates and preferred habitats^11^. In the wild, it thrives in complex moist and liquiform environments such as soil and rotten fruits^12^. These surroundings present challenges and require quick, energy-efficient decisions to survive and compete for food or mating partners. *C. elegans* belongs to the animal class *Secernentea*, whose members feature sensilla (i.e., sensory apparatuses) in their anterior (head) and posterior (tail) sides (amphids and phasmids, respectively)^13–17^. One cell-type associated with the amphids are the ASH neurons, which are polymodal nociceptive neurons that can detect a wide range of aversive stimuli^18^. ASHs play key roles in stimulus-evoked backward locomotion, mostly referred to as escape behavior^19,20^ (Figure 1A). The phasmids feature the ciliated sensory neurons PHA and PHB, located in the tail’s sensory organs; these neurons have been shown to act as polymodal sensory neurons that sense harmful chemicals, hyperosmotic solutions and mechanical stimulation^21^ (Figure 1A). These head and tail neurons share many similarities, such as neuronal activity patterns and polymodality for noxious cues^21,22^. While ablation of ASH neurons causes a significant reduction in avoidance responses, following the additional ablation of PHA and PHB, the behavioral response returns to normal^19^. This finding strongly suggests that PHA and PHB negatively modulate the escape response. However, the mechanism underlying such modulation has yet to be elucidated.

**Figure 1.**
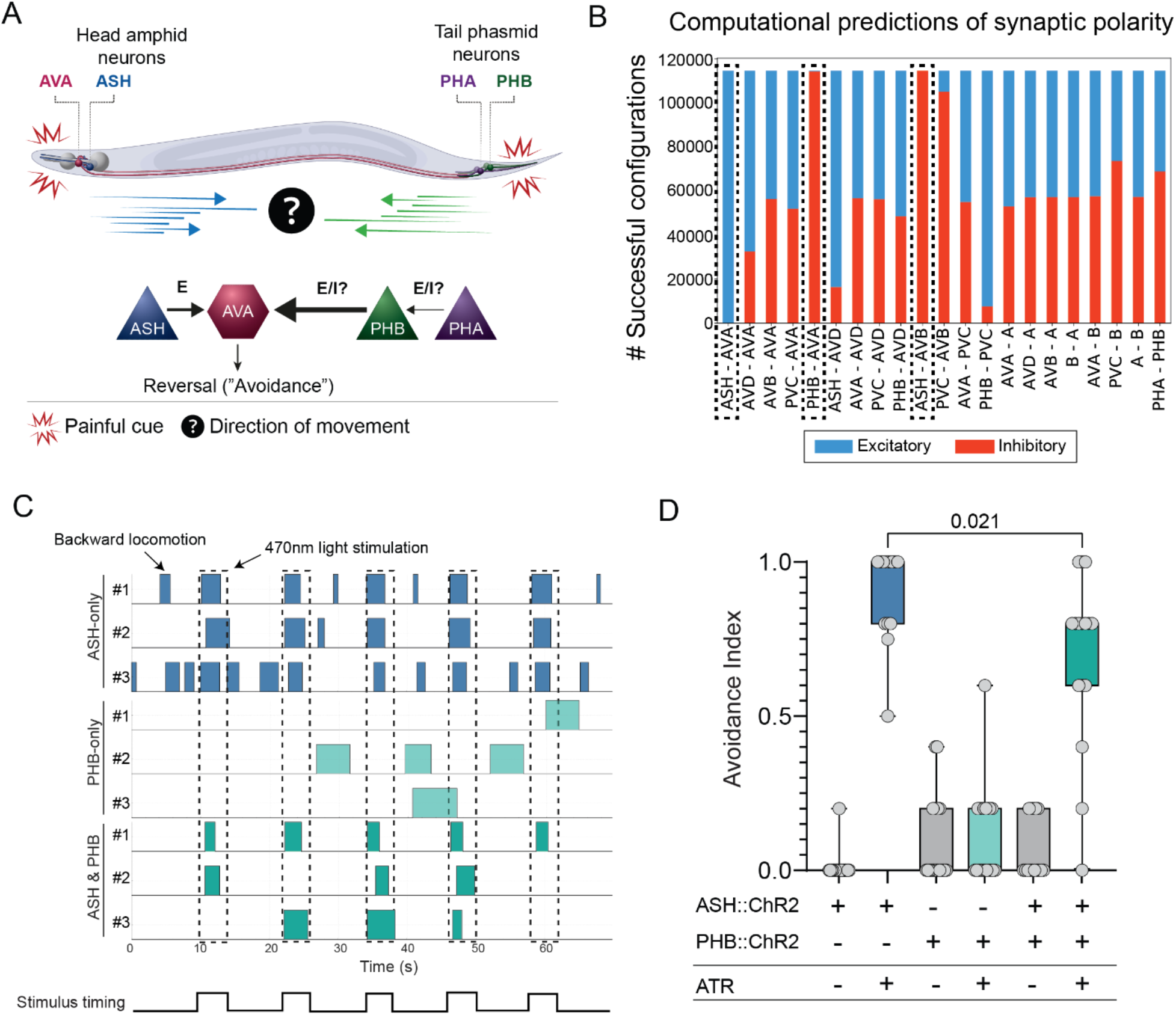
Head and tail stimulations result in antagonistic behaviors. (A) Schematic illustration of the application of a painful stimulus to the head and tail of *C. elegans*. Arrows depict backward locomotion following head stimulation (blue) and forward locomotion following tail stimulation (green). E, excitation; I, inhibition. (B) Computational predictions of synaptic polarity in the circuit for nociceptive behaviors. All the combinations of synaptic polarity were tested and defined as successful based on several conditions (see Methods), including the suppression of backward locomotion by the activation of tail sensory neurons (PHA, PHB). Most synapses can be either excitatory (blue) or inhibitory (red), whereas only three connections were strictly excitatory (ASH>AVA) or inhibitory (PHB>AVA, ASH>AVB) across over 99% of successful configurations (dashed-line boxes). See Methods for a full description of the model. (C) Representative avoidance responses of individual worms to five consecutive optogenetic activations (blue-light stimuli, dashed lines). *ASHp∷ChR2* (ASH-only), *PHBp∷ChR2* (PHB-only), *ASHp∷ChR2;PHBp∷ChR2* (ASH & PHB). Plotted boxes represent reversal events. (D) The avoidance index for sensory neuron activation. Evoked reversal responses to ASH stimulation are suppressed by the simultaneous stimulation of the PHB neuron. The avoidance index was calculated as the fraction of reversal responses from a total of five stimulations of each individual. n=13–20. In D, we performed Mann-Whitney test.

Electron microscopy reconstructions of the nervous system of *C. elegans* indicate that only two neurons are innervated by both ASH and PHB – interneurons AVA and AVD^23,24^ – implying that they may be hubs of spatial information computation. Both AVA and AVD mediate the initiation of backward locomotion (reversal behavior), with AVA shown to play the major role ^25,26^. AVA is active during reversals and in response to signaling from ASH^27,28^. The glutamatergic identities of ASH and PHB suggest that AVA receives simultaneous excitatory and inhibitory glutamate-mediated inputs from these spatially distinct neurons (Figure 1A)^27,29^. Nevertheless, how AVA computes spatial information, integrates the signals and produces beneficial behaviors remains unknown.

While glutamate neurotransmission activates AVA through different types of AMPA- and NMDA-like receptors, glutamate-gated chloride channels (GluCls) have been suggested to mediate weak inhibitory currents^30,31^. Such integration of excitatory and inhibitory signals in single neurons is associated with sensorimotor circuits that are responsible for computation of multisensory cues^32,33^. In *C. elegans*, different GluCls are active in specific neurons and modify various behaviors such as ivermectin sensitivity, salt chemotaxis, thermotaxis and spontaneous reversal rate^34–37^.

Here, we demonstrate that the architecture of a neuronal circuit can serve as the infrastructure for sensing and integrating multidimensional information, such as the location and concentration of stimuli in the environment. We show that activation of tail sensory neurons suppresses *C. elegans*’s avoidance behavior induced by head sensory neuron activation. By recording the neuronal activity of ASH, PHA, PHB and AVA following exposure of the head or tail to noxious stimulus, we reveal that while head stimulations result in both ASH and AVA activation, tail stimulations elicit activation in PHA and PHB but inhibition in AVA, in a concentration-dependent manner. We examined AVA’s response to stimulus under mutant glutamate receptor backgrounds and found that head-evoked excitation in AVA depends on the excitatory GLR-1 and NMR-1. In contrast, tail-evoked inhibition requires the inhibitory AVR-14, specifically in AVA. The imaging of GFP-tagged receptors revealed a spatially partitioned localization of the excitatory and inhibitory channels along the AVA process. Finally, using behavioral assays and optogenetics to analyze stimulus-evoked reversal rates in mutant backgrounds, we found that GLR-1/NMR-1-mediated neurotransmission is crucial for normal avoidance responses, and that AVR-14 suppresses the reversal rate in an AVA-specific manner. Taken together, our findings describe the neuronal and molecular mechanisms that converge onto a single interneuron, which then relays the decision whether or not to initiate an avoidance behavior.

## RESULTS

### Antagonistic functions of head and tail signaling

Previous work indicated that ASH is the main nociceptive neuron, functioning through a compact circuit that controls nociceptive behaviors^19,28^ (Figure 1A). Using a computational model we previously developed, we predicted which connections in the circuit could be inhibitory^28^. To do so, we added PHA and PHB to our previous circuit simulation and tested all the combinations of excitatory and inhibitory connections, such that each connection can be either inhibitory or excitatory (see Methods section for details). Our model predicted that most connections can be either excitatory or inhibitory and still maintain the proper function of the network (Figure 1B). However, few connections had a much higher tendency for a certain polarity: ASH>AVA was almost exclusively excitatory, whereas PHB>AVA and ASH>AVB almost exclusively inhibitory. Thus, our model is able to predict behavioral outcomes of the integration of excitatory and inhibitory connections.

To examine whether PHB>AVA is indeed inhibitory and, more broadly, determine the contribution of specific sensory neurons in the circuit to avoidance behavior in freely moving animals, we utilized an optogenetic approach. We photostimulated ASH, PHB or both simultaneously in transgenic animals expressing Channelrhodopsin-2 (ChR2) under cell-specific drivers, and then analyzed their avoidance behavior. While ASH-specific photo-activation induced strong avoidance responses, as previously reported^28,38^, PHB-specific stimulation did not evoke any avoidance behavior (Figure 1C-D). However, the combined activation of both ASH and PHB elicited a significantly reduced avoidance response (Figure 1C-D). Since AVA is the main backward command interneuron connected to ASH and PHB, these results suggest that the PHB>AVA synapse is inhibitory and attenuates the avoidance response initiated by an excitatory input from the ASH>AVA synapse.

### Head or tail exposure to a nociceptive cue induces distinct activity patterns

The sensory neurons ASH, PHB and PHA become active when their cilia are exposed to a variety of aversive chemicals^21,22,28,38,39^. Although the chemical selectivity of the neurons is well established, their sensitivity to different concentrations of chemicals has not been defined, despite evidence of dose-dependent behavioral responses to some aversive cues^28^. To compare the sensitivity of the sensory neurons, we recorded the calcium level changes in ASH (‘head’), and PHB and PHA (‘tail’), following exposure to the osmotic stressor glycerol at two different concentrations (0.5 M and 2 M). We used the ‘olfactory chip’^40^, a microfluidic device, to image GCaMP6s-expressing animals with either their head or tail exposed to the stimulus. ASH was activated following head exposure to both glycerol concentrations (Figure 2A). However, in both PHA and PHB, tail exposure to glycerol induced calcium activity only in response to the higher concentration (Fig. 2B, C). Furthermore, ASH and PHA showed an additional OFF response upon stimulus removal (Fig. 2A, C). Thus, head ASH and tail PHA and PHB sensory neurons show distinct activity patterns and activation thresholds.

**Figure 2.**
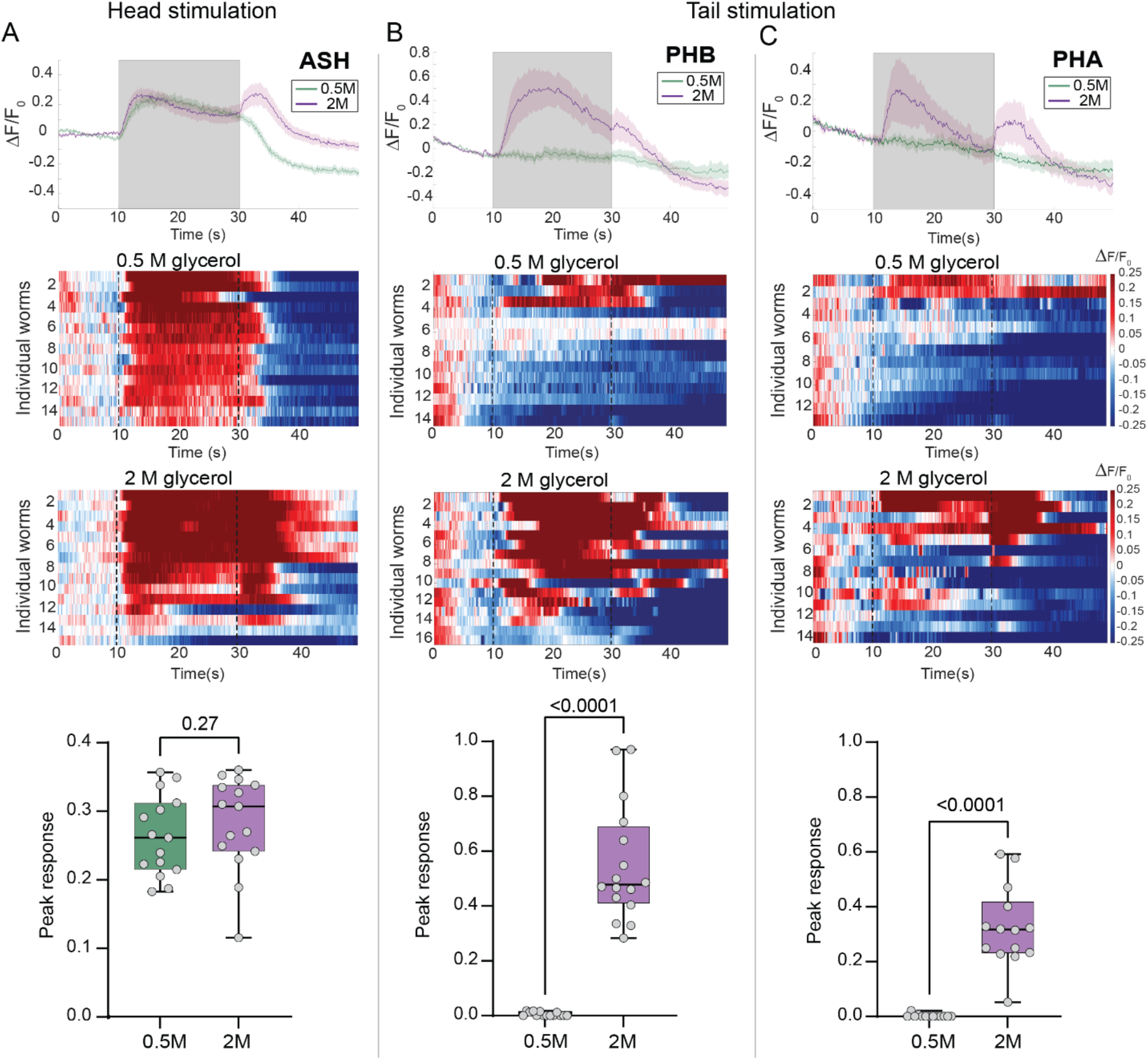
Head or tail exposure to a nociceptive cue induces diverse activity patterns of sensory neurons. (A-C) Calcium traces in the ASH sensory neuron (A) and PHB and PHA neurons (B-C) following head and tail stimulation, respectively, with 0.5 M or 2 M glycerol. Top, average and SEM traces of calcium responses to 0.5 M glycerol (green) and 2 M glycerol (purple). Gray background indicates the time of stimulus delivery. Middle, normalized, color-coded GCaMP6s calcium responses of individual animals exposed to each concentration. Heatmaps represent the calcium levels in individual worms. The stimulus was applied at 10–30 s. Bottom, quantification of peak responses (see Methods). In A-C, we performed Mann-Whitney tests. n=13–16 animals per group.

### AVA interneuron is activated by head stimulation and inhibited by tail stimulation

Given that AVA is innervated by the sensory neurons, this neuron may be equipped with a mechanism for analyzing the location and concentration of stimuli it receives. For this reason, we next turned to identify the mechanism by which AVA simultaneously integrates spatially conflicting neuronal signals that represent different threshold sensitivities.

Although it was previously shown that AVA becomes activated following chemical head stimulation^28^, its activity patterns following tail stimulation have not been reported so far. We, therefore, assessed the changes in calcium levels in AVA in response to head or tail stimulation with high and low glycerol concentrations. Whereas stimulating the head with both tested concentrations induced activity in AVA (Figure 3A), only the application of the high concentration in the tail induced inhibition in AVA (Figure 3B).

**Figure 3.**
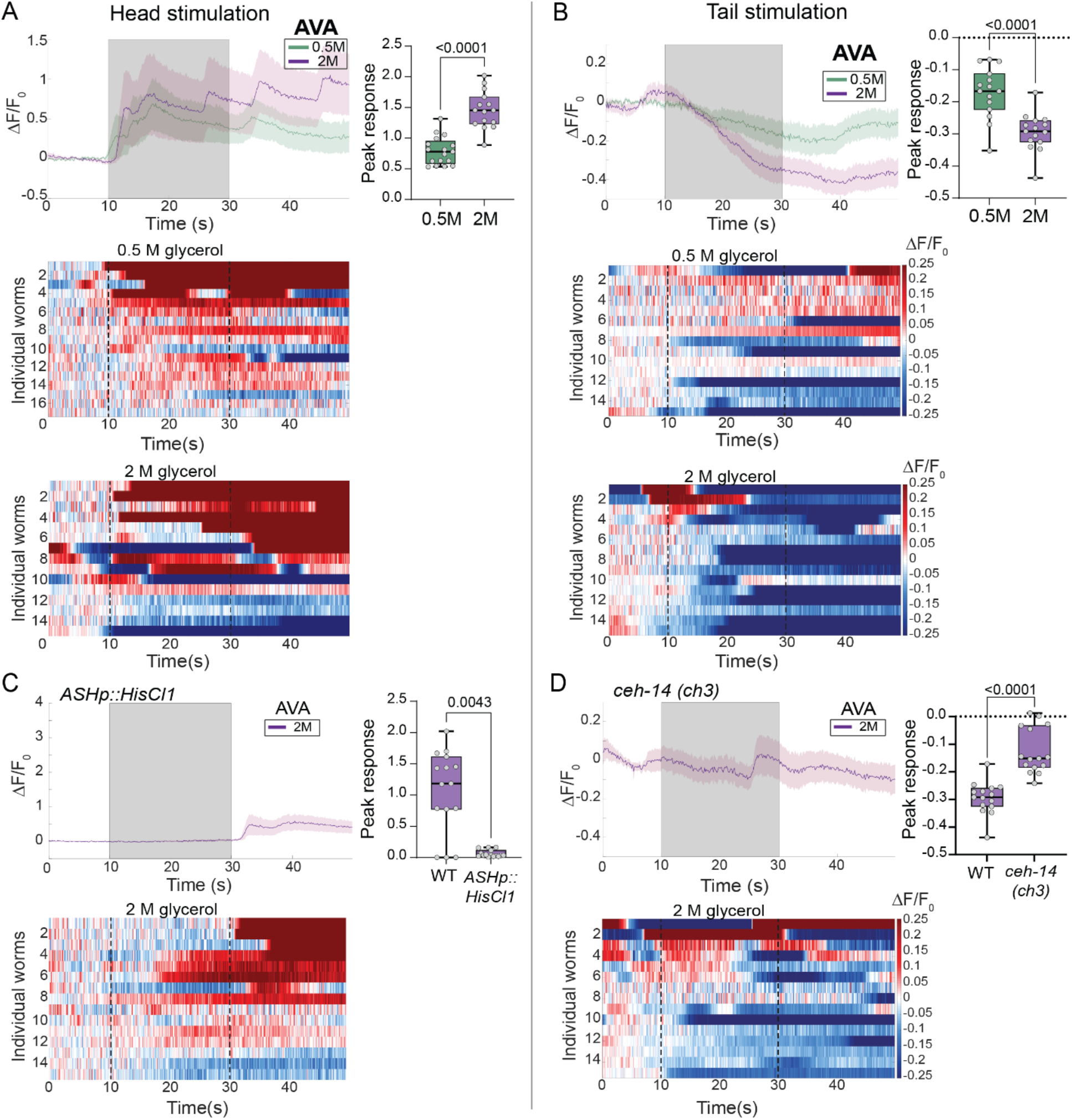
AVA interneuron is activated following head exposure and inhibited following tail exposure to nociceptive stimuli. (A-B) GCaMP6s calcium responses in AVA following head (A) or tail (B) stimulation with 0.5 M or 2 M glycerol. Top, average and SEM traces, and quantification of peak responses. Bottom, heatmaps of the normalized calcium levels (color-coded) of individual animals. Stimulus was applied at 10–30 s. (C) AVA calcium responses in ASH-silenced animals following head stimulation with 0.5 M or 2 M glycerol (see Methods). Top, average and SEM traces, and quantification of peak responses. Bottom, heatmaps of the normalized calcium levels (color-coded) of individual animals. (D) AVA calcium responses in *ceh-14* mutant animals following tail stimulation with 0.5 M or 2 M glycerol. Top, average and SEM traces, and quantification of peak responses. Bottom, heatmaps of the normalized calcium levels (color-coded) of individual animals. n=15 animals per group. In A-D, we performed Mann-Whitney tests.

To test whether the head and tail sensory neurons are essential for AVA activation, we measured the neuronal activity following strong stimulation (2M glycerol) in animals with impaired head or tail sensory neurons. We silenced ASH using cell-specific expression of the inhibitory *Drosophila* histamine-gated chloride channel (HisCl1)^41^, and analyzed mutants of the *ceh-14/Lhx3* LIM homeobox gene, in which glutamate transmission from PHA and PHB is defective^29^. We found that ASH-silenced animals show significantly weaker activation in AVA following head stimulation in comparison to wildtype animals (Figure 3C). In contrast, *ceh-14* animals showed no inhibition following tail stimulation (Figure 3D). Taken together, our data show that the head and tail sensory neurons mediate excitatory and inhibitory currents in AVA, respectively, in response to noxious stimuli.

### Simultaneous head and tail stimulation evoke AVA activity in a concentration-dependent manner

In the natural environment of the worm, both its head and tail are exposed to stimuli. Therefore, to decipher the neuronal activity dynamics in AVA in a more naturalistic environment, we designed and fabricated a new microfluidic device, the ‘dual olfactory chip’, which allowed us to stably position worms with both their head and tail exposed to a buffer flow (Figure 4A-B, see Methods). We then used this chip to visualize AVA calcium currents following simultaneous chemical stimulation of the head and tail. Remarkably, concurrent head and tail stimulation of AVA induced a distinct activation pattern (Figure 4C-D) that differed from that induced by either head or tail stimulation alone (Figure 3): AVA became active only following low-concentration chemical stimulation. This finding implies that such a stimulus suffices to induce activity in AVA but not to pass the tail-activation threshold needed to induce its inhibition. Thus, tail sensation antagonizes head-induced activity at the interneuron level, which leads to the production of an integrated dosage-dependent response to an aversive cue (Figure 4E).

**Figure 4:**
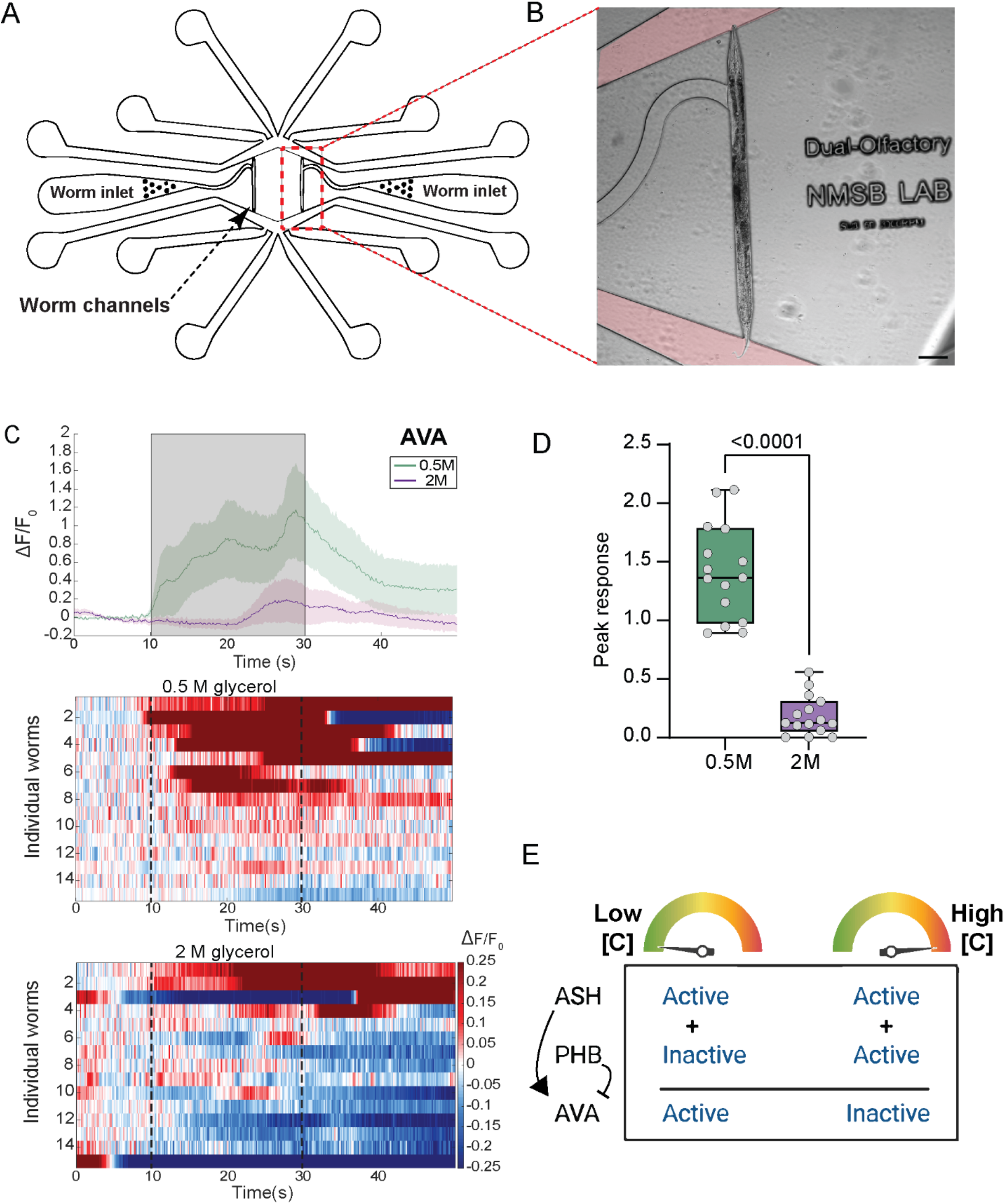
Simultaneous head and tail exposure to nociceptive stimuli reveals a concentration-dependent activity pattern in AVA. (A) Schematics of the ‘dual olfactory chip’. Worms are inserted to the chip through the worm inlet and trapped in the worm channel (dashed-line box). The buffer and stimulus flow into the chip through 8 inlets (4 inlets at the top and 4 at the bottom). The stimulus flow direction is controlled using a manual valve. The fluid flows out of the chip through 4 outlets (see Methods for full details). (B) Representative image of a worm trapped in the worm channel of the ‘dual olfactory chip’. Red dye, stimulus flow. Scale bar, 100 μm. (C) Calcium traces in AVA following the simultaneous stimulation of the head and tail with 0.5 M (green) or 2 M (purple) glycerol. Top, average and SEM traces. Bottom, heatmaps of the normalized calcium levels (color-coded) of individual animals. Stimulus was applied at 10–30 s. (D) Quantification of peak responses. n=15 animals per group. We performed a Mann-Whitney test. (E) Schematic model describing how the different activation thresholds of the sensory neurons in the head (ASH) and tail (PHB) can affect the activity pattern of downstream interneuron AVA. The computation of concentration-independent excitation from the head and concentration-dependent inhibition from the tail results in either the activation or inactivation of AVA, probably affecting behavioral responses to aversive cues.

### Distinct excitatory and inhibitory glutamate-gated receptors mediate AVA activity

Since glutamatergic signals from PHB to AVA are required for AVA inhibition following tail stimulation, we sought to identify the glutamate receptors expressed in AVA that can mediate these inhibitory currents. Two *inhibitory* glutamate-gated chloride channels (GluCls) have been reported to be expressed in AVA: AVR-14 and GLC-4^31,34,36,42,43^(Figure 5A). We analyzed the calcium levels in AVA following tail stimulation in *avr-14* and *glc-4* mutant animals, and found that while AVA inhibitory currents remained intact in *glc-4* mutants, they were completely abolished in *avr-14* mutants (Figure 5B,C,F). In contrast, head stimulation of *avr-14* mutants did not alter AVA activity (Figure 5G,I). Expressing AVR-14 specifically in AVA under an *avr-14* mutant background sufficed to rescue AVA inhibition following tail stimulation (Figure 5D,F). Thus, *avr-14* is required specifically in AVA to mediate inhibitory currents following tail stimulation.

**Figure 5.**
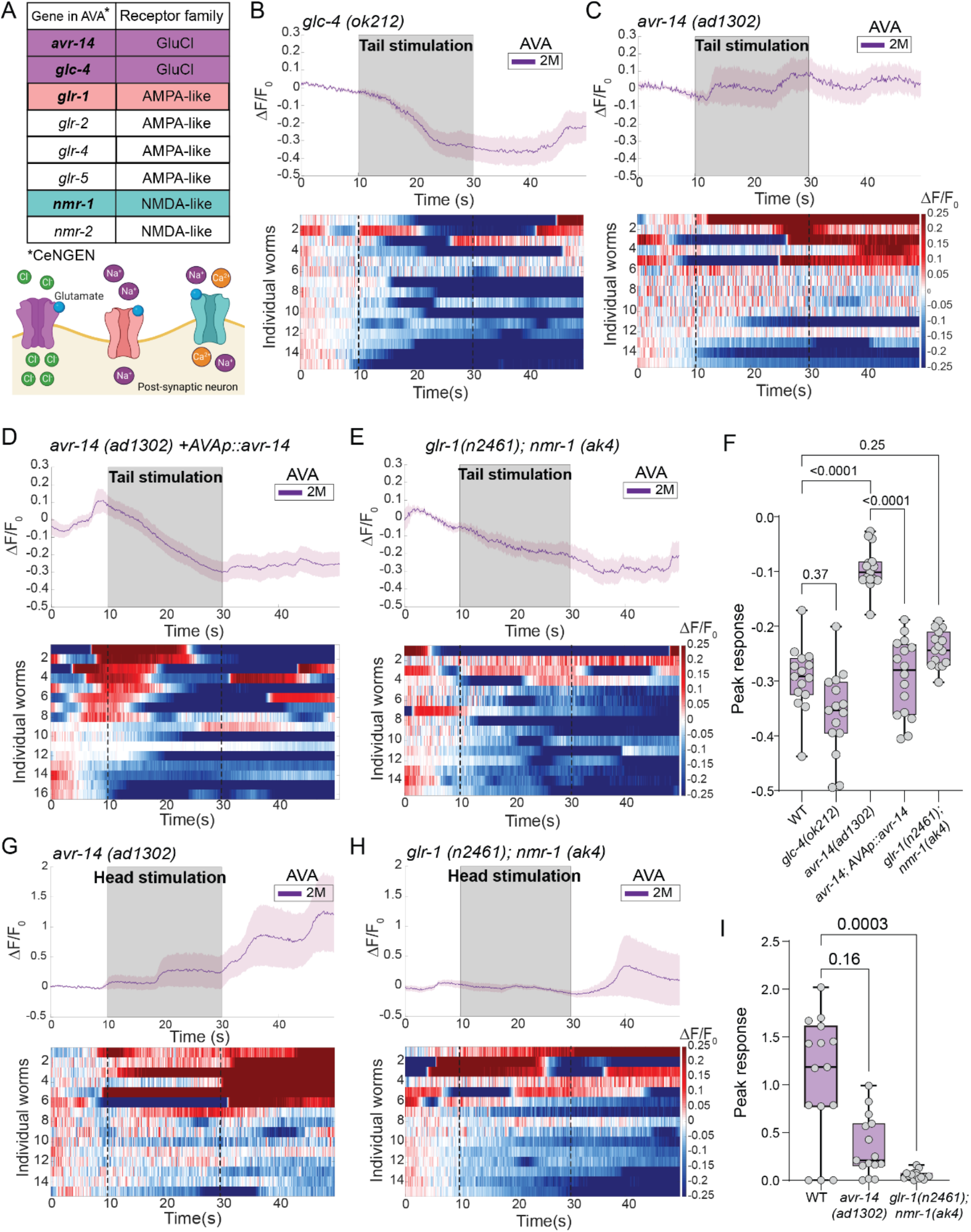
Various excitatory and inhibitory glutamate-gated receptors control activity dynamics in AVA. (A) Top, glutamate receptors encoding genes predicted to be expressed in AVA^43^ and their families. The two types of GluCl inhibitory receptors, *glc-4* and *avr-14* (purple), the AMPA-like *glr-*1 (red) and the NMDA-like *nmr-1* (turquoise) excitatory receptors, were selected to study neuronal activity dynamics in AVA upon head or tail stimulation. Bottom, schematic illustration of the various types of glutamate receptors and their excitation/inhibition conductance mechanisms. (B-E) AVA calcium responses following tail stimulation with 2 M glycerol in *glc-4* mutants (B), *avr-14* mutants (C), *avr-14* mutants expressing *avr-14* specifically in AVA (D), and *glr-1, nmr-1* double mutants (E). Top, average and SEM traces. Bottom, heatmaps of the normalized calcium levels (color-coded) in individual animals. Stimulus was applied at 10–30 s. (F) Quantification of peak responses. (G-H) AVA calcium responses following head stimulation with 2 M glycerol in *avr-14* mutants (G) and *glr-1, nmr-1* double mutants (H). Top, average and SEM traces. Bottom, heatmaps of the normalized calcium levels (color-coded) in individual animals. Stimulus was applied at 10–30 s. (I) Quantification of peak responses. n=15–16 animals per group. In F, we a performed Kruskal-Wallis test followed by a Dunn’s multiple comparison analysis.

Although various *excitatory* glutamate receptors are expressed in AVA (Figure 5A)^43^, we focused on the GLR-1 AMPA-like and NMR-1 NMDA-like receptors, as they were shown to play a key role in AVA-evoked excitatory currents^27,31,44^. We analyzed neuronal activity in the AVA of *glr-1;nmr-1* double mutants following head or tail high-concentration stimulation. While these double mutants showed almost no AVA activation following head stimulation, AVA inhibition following tail stimulation was intact (Figure 5E,F,H,I). Taken together, our data might indicate that AVR-14, GLR-1 and NMR-1 are unevenly localized post-synaptically along the AVA process.

### Differential localization of distinct glutamate receptors along the AVA process

Previous work has shown that differential cellular localization of distinct types of glutamate receptors can mediate the integration of excitatory and inhibitory currents, to fine tune behavior^37,45,46^. To analyze the localization of glutamate receptors along the AVA process, we generated transgenic animals expressing fluorescent-tagged GLR-1, NMR-1 and AVR-14 proteins in AVA (see Methods) and quantified the number of puncta in four distinct regions along its process (Figure 6A). Although puncta of all three receptors were visible in all regions, GLR-1 and NMR-1 were enriched in anterior regions of the AVA process, while AVR-14 was enriched posteriorly (Figure 6B-E). The trafficking of synaptic proteins (including GLR-1) in AVA was previously shown to be mediated by kinesin-3 UNC-104(KIF1A)^47–50^. To determine whether *unc-104* is involved also in AVR-14 trafficking and localization, we analyzed the AVR-14 puncta density and soma expression in AVA in *unc-104* mutant animals (Figure S1A). Although the mean intensity levels in AVA soma were similar in wildtype and *unc-104* animals, we found absolutely no puncta along the AVA process in an *unc-104* mutant background (Figure S1B-D), suggesting that UNC-104 is required for AVR-14 trafficking to the posterior regions of the AVA process. Taken together, our results points to the differential localization of inhibitory and excitatory receptors along the AVA process as one mechanism through which AVA encodes spatially conflicting cues to possibly drive avoidance behavior.

**Figure 6.**
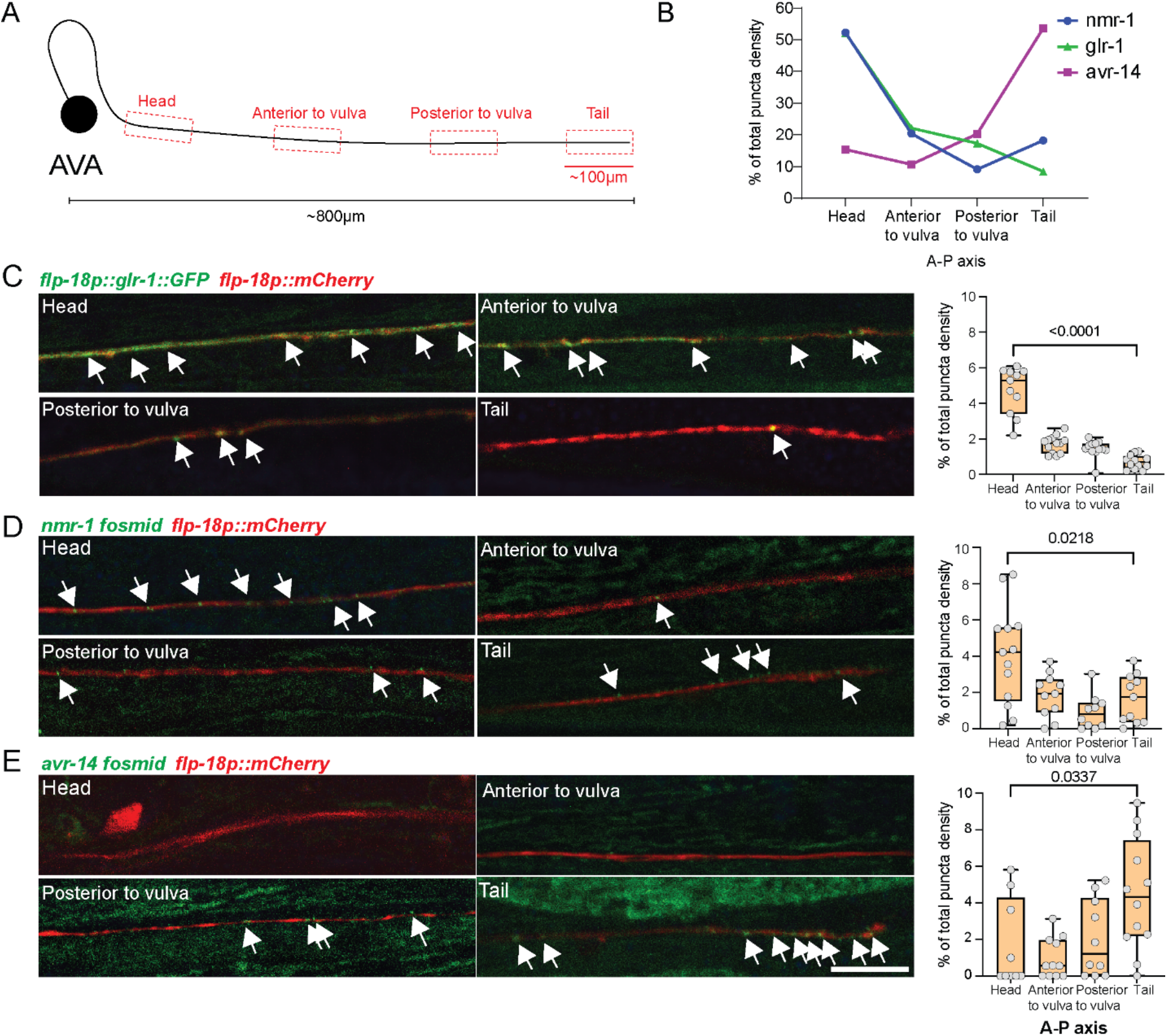
Distinct types of glutamate receptors are localized differentially along the AVA process. (A) Schematic of the regions in the AVA process that were analyzed (dashed-line red boxes). Each analyzed region along the process is ~100 μm in length. The AVA process’s length is roughly ~800 μm. (B) Quantification of NMR-1 (blue), GLR-1 (green) and AVR-14 (purple) puncta localization along the AVA process. Each dot represents the proportion of GFP puncta in each region along the anterior-posterior axis. GLR-1 and NMR-1 localize mostly to the anterior regions (>70%), while AVR-14 localizes mostly to the posterior regions (>70%). (C-E) Representative confocal images and quantifications for all four regions of the AVA process quantified in (B) in animals expressing a red fluorescence marker in AVA (*flp-18p∷mCherry*) along with GFP-fused reporters of the glutamate receptors *flp-18p∷glr-1∷gfp* (C), *nmr-1 fosmid* (D), and *avr-14 fosmid* (E). White arrows denote puncta locations. n=9-13 animals per group. In C-E we performed a Mann-Whitney test.

### Avoidance behavior is modulated by the concerted action of GLR-1, NMR-1 and AVR-14

To analyze the contribution of the different glutamate receptors to the avoidance behavior, we used the drop test^19,20,28^ (see Methods). In this test, both the head’s and tail’s sensory neurons are stimulated by applying a drop of either a high- or low-concentration of glycerol to forward-moving animals and then their reversal rates are scored. We found that while *avr-14* mutants exhibited normal avoidance behavior, the responsiveness of *glr-1;nmr-1* double mutants was almost completely abolished, in a manner independent of stimulus concentration (Figure 7A). These results imply that AVA fails to become active and evoke a behavioral response in the absence of excitatory, but not inhibitory, glutamate receptors. The *glr-1;nmr-*1*;avr-14* triple mutants only showed a significantly higher avoidance index than the *glr-1;nmr-*1 double mutants in response to the high concentration (Figure 7A). This finding suggests there is some residual GLR-1/NMR-1-independent activation mechanism at play in AVA, which could be GLR-2-mediated^31^. The AVA-specific rescue of *avr-14* expression in the triple-mutant background was sufficient to significantly decrease the avoidance behavior in animals that were exposed to the high-concentration stimulus (Figure 7A), indicating that *avr-14* is necessary specifically in AVA to mediate the repression of avoidance behavior.

**Figure 7:**
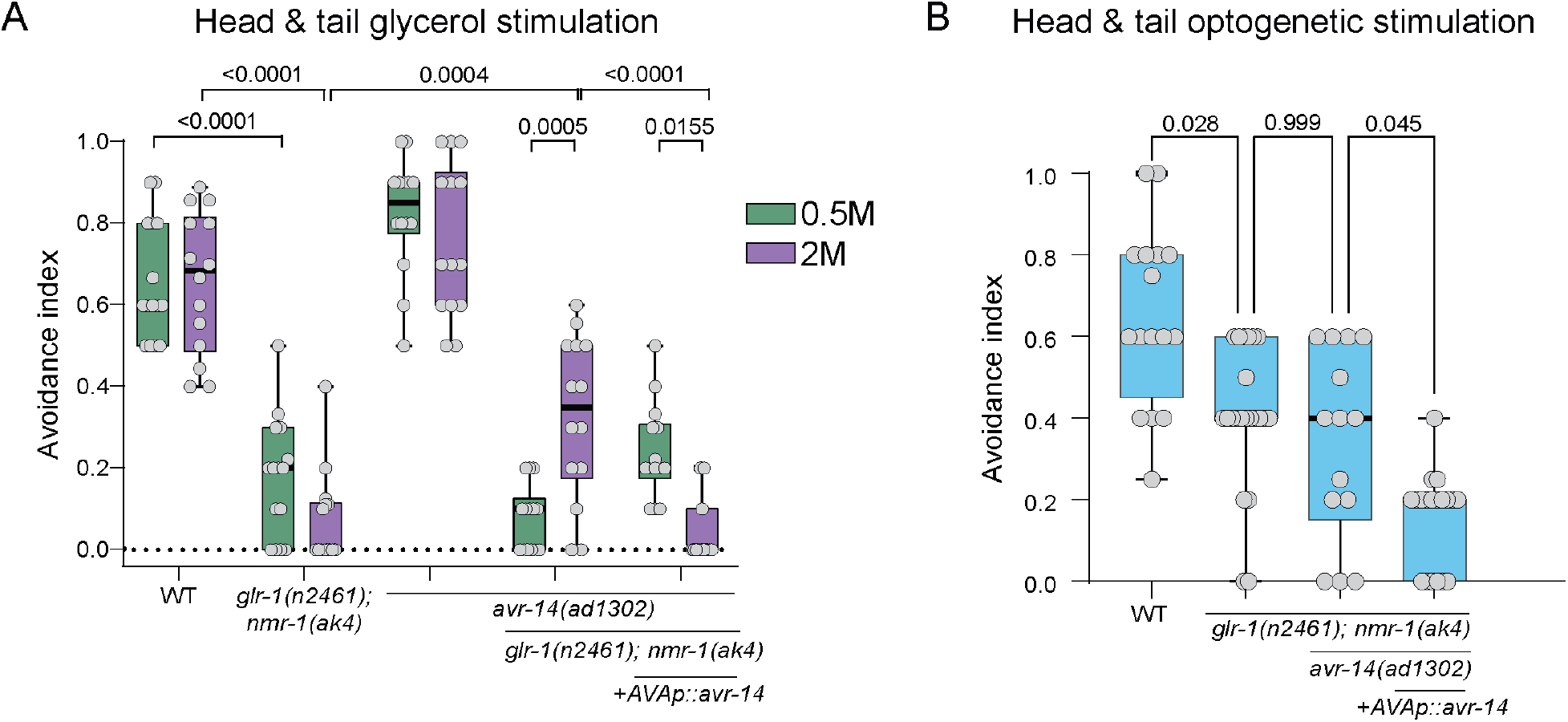
Behavioral decision is regulated by GLR-1, NMR-1 and AVR-14 receptors. (A) Behavioral responses to 0.5 M (green) or 2 M (purple) glycerol using the drop assay (see Methods). The avoidance index represents the fraction of reversal responses in 8-14 trials of each single animal. n=14–15 worms per group. (B) Avoidance behavior in response to the simultaneous optogenetic stimulation of both ASH and PHB. n=14–22 animals per group. In (A), we performed Two-way ANOVA, Tukey’s multiple comparisons test. In (B), we performed One-way ANOVA, Kruskal-Wallis test followed by Dunn’s multiple comparison analysis.

As a complementary approach, we used optogenetics to photo-activate ASH and PHB neurons and measured the evoked avoidance responses in different mutant backgrounds. This approach allowed us to determine ASH- and PHB-specific contributions to behavior, by avoiding the stimulation of other sensory neurons and bypassing the tail-activation threshold. The *glr-1;nmr-1* double mutants and *glr-1;nmr-1;avr-14* triple mutants showed a similar decrease in avoidance response compared to wild-type animals (Figure 7B). Interestingly, the decrease in the *glr-1;nmr-1* avoidance response was only partial, unlike the complete loss we observed in the drop assay, suggesting that the optogenetic stimulation setup does not fully mimic chemical stimulation. One possible reason for this could be the uneven expression of ChR2 in ASH and PHB. As expected, AVA-specific expression of *avr-14* in the *glr-1;nmr-1;avr-14* mutant background was sufficient to completely abolish the avoidance behavior (Figure 7B). Taken together, these results indicate that AVA-specific activity of AVR-14 suppresses GLR-1/NMR-1 mediated avoidance responses, and that ASH>AVA and PHB>AVA are major but not the sole signaling pathways orchestrating this behavior.

## DISCUSSION

Here we show how *C. elegans* utilizes a compact neuronal circuit to recognize hazards in a complex environment, integrate spatial information and make a decision. We find that this process is supervised by at least three properties: (i) the properties of the sensory neurons, i.e., spatial receptive field, sensitivity and anatomical location, (ii) the properties of the environmental stimulus, i.e., location and concentration, and (iii) the localization patterns of excitatory and inhibitory channels along the process of the interneuron. Although not addressed here, neuromodulation might also play a role.

Avoidance behavior in worms is an evoked response, characterized by backward locomotion followed by a turn to change the direction of movement^51^. We examined the contribution of the major backward command interneuron AVA to the decision-making process of the avoidance behavior and found that it plays a key role in integrating multiple distinct signals in order to generate or abort the initiation of backward locomotion. We found AVA to be inactivated when the head and tail were strongly stimulated, but behaviorally, animals did show high avoidance responses, pointing to a redundant mechanism in which AVA is a key but not the sole mediating neuron of avoidance behavior. Other backward command interneurons such as AVD (which is innervated by both head ASH and tail PHB sensory neurons) and AVE might also act to guide avoidance behavior^25^. Furthermore, the AIB-RIM disinhibitory motor circuit, which was shown to be stimulated by ASH and affect AVA activity, may contribute to the reversal decision^27,52^. The AVA interneurons have also been suggested to function as hub neurons which integrate sensory inputs from threat and reward circuits along with motor information and guide decision making^53^, but whether this occurs in a spatial manner remains to be determined.

We further describe the molecular mechanism that is utilized to integrate spatially opposing stimuli within a single neuron. We show that glutamate signaling can cause excitation or inhibition in AVA, leading to an increase or a decrease in the probability of reversal, respectively, and that these neuronal responses depend on the receptor type. We also demonstrate how differential localization of excitatory (i.e., AMPA- and NMDA-like) and inhibitory (GluCl) receptors along the interneuron process dictate the opposing neuronal responses following head and tail stimulations. Although we tested the role of significant glutamate receptors, such as the excitatory GLR-1 and NMR-1 or the inhibitory GLC-4 and AVR-14, other receptors may play a role. Possible candidates include GLR-2, known to participate in AVA activation, and GLR-4/5 and NMR-2, which are expressed in AVA^31,43^. Evidence suggests that at least some of these glutamate receptors, such as GLC-4 and AVR-14 or GLR-1 and NMR-1, work in concert to modulate their activity^21,31^. Whether the singular effect of each receptor or the combinatorial summation are important for avoidance decisions remains an open question.

Since glutamate secretion from the head ASH and tail PHB sensory cells can be evoked by various stimuli, we predict that different modalities other than osmo-sensation, such as response to touch, toxins and other repellents, will show similar neuronal dynamics and behavioral modulation following exposure to spatially opposing cues^19,21,22,28,52,54,55^.

GluCls are found only in protostome invertebrate phyla, but are closely related to mammalian glycine receptors^56^. In mammals, Glycine receptor Cl- channels (GlyRs) play an important role in rapid synaptic inhibition in the spinal cord, brainstem and higher brain centers, and are involved in the transmission of nociceptive signals^57^. Recently, alpha3-subunit-containing GlyRs have been implicated in the inflammation-mediated disinhibition of centrally projecting nociceptive neurons, marking them as novel molecular targets in pain therapy^58,59^. Taken together, these findings highlight the importance of understanding the mechanisms of action of these channels for future pain-related therapeutic approaches.

ASH, PHA and PHB sensory neurons show diverse activity patterns following exposure to noxious stimuli at different concentrations. We uncovered a differential activation threshold: while the head ASH neurons are activated in response to low and high concentrations, the tail PHA and PHB neurons respond only to high concentrations. The different sensitivity may result from an asymmetry in the expression of molecular elements that govern neuronal sensation, such as TRPV channels OSM-9/OCR-2, various types of GPCRs or OSM-10^19,43,60,61^. Additionally, ASH neurons show an OFF response only at a high concentration, implying that it may mediate some concentration-dependent behaviors. Finally, we discovered that the PHA neuron, which innervates PHB and AVG, but not AVA (in hermaphrodites), displays concentration-dependent ON and OFF responses, which correspond to neuronal responses in AVG following tail exposure to touch and high-concentration osmotic shock^62,63^. This finding suggests an additional parallel pathway, mediated by PHA>AVG, that respond to osmotic shock applied to the tail, but its contribution to behavioral decisions remain unclear. The significance of slight differences in the location of stimuli in the surroundings is well demonstrated by the tendency of *C. elegans* to suppress exploratory head movements only following light anterior touch, as opposed to nose and posterior touch, in a mechanism thought to enable escape from traps of predatory fungi^61^. Taken together, our results demonstrate how a relatively simple neural network can encode multidimensional information, including the spatial location and concentration of stimuli. Despite the small number of neurons and the single neurotransmitter (glutamate), the circuit utilizes distinct types of receptors, which are differentially localized, to integrate information and quickly direct behavioral decisions.

## METHODS

### Experimental Model and Subject Details

All *Caenorhabditis elegans* strains that were used in this study are listed in supplementary table 1. Worms were maintained according to standard conditions, at 20°C on nematode growth medium (NGM) plates that were seeded with *E. coli* OP50 bacteria^64^. The Bristol N2 strain was used as wild-type control. *him-5*(e1490) were treated as wild-type controls for strains carrying this allele in their genetic background.

## Method details

### Molecular biology

To generate the *avr-14* rescue construct, pAG1, and capture all the *avr-14* isoforms, we fused a cDNA fragment encoding the first 6 common exons with a genomic fragment containing the rest of the gene, as previously described^27^. This fragment was cloned under the flp-18 promoter for AVA-specific expression.

To generate the specific expression of GLR-1∷GFP in AVA, we attached the GLR-1∷GFP construct^65^ to the *flp-18* promoter, to generate pAG4.

To express ChR2 in PHB neurons, we cloned a *ChR2∷mCherry* fusion under the PHB-specific *gpa-6* promoter, to generate pHS9.

### Confocal microscopy

Animals were mounted on a 5 % agarose pad on a glass slide, on a drop of M9 containing 100– 200 mM sodium azide (NaN3). A Zeiss LSM 880 confocal microscope was used with 63x magnification. For *nmr-1/avr-14* fosmids, AVA was identified using *flp-18p∷mCherry*, and the z-plane with the strongest signal was chosen. Puncta densities were calculated by normalizing the number of puncta to the length of the relevant part of the AVA process (regions from anterior to posterior: head, anterior to vulva, posterior to vulva, tail). For the *GLR-1∷GFP* analysis, the signal was mostly very strong and puncta were difficult to separate. We measured the length of an average puncta using a sample of 3–5 puncta from each animal, and analyzed puncta density by dividing the length of the relevant process by the average length of the puncta. The fluorescence intensity in the soma was analyzed using ImageJ version 1.52p. The relevant Z-slices were sub-stacked, “Sum slices” was applied, and Regions of interest (ROIs) were selected before the measurement and export of the mean intensity.

### Histamine-induced silencing

Histamine plates were prepared as previously described^41^. NGM-histamine (10 mM) and control plates were stored at 4°C for no longer than 2 months. Histamine plates were tested using worms that carry a transgene with a pan-neural HisCl1 (*tag-168∷HisCl1∷SL2∷GFP*)^41^. After a few minutes on histamine plates, these worms were paralyzed completely, validating the potency of the histamine plates.

### Microfluidic chip fabrication and calcium imaging

#### Olfactory chip fabrication

The olfactory chip was fabricated according to^40^ with the help of the Nanofabrication Unit at the Weizmann Institute of Science. A polydimethylsiloxane (PDMS) mixture was cast into premade 0.5-cm-high chip molds and allowed to solidify at 65°C for 3 h. Individual chips were cut by hand with a scalpel and then punctured to create fluidic inlets with a diameter of 0.5 mm, using a PDMS biopsy punch (Elveflow). The chips were attached to glass coverslips by exposing them to plasma for 30 s, and then manually attaching them together and drying them on a hot plate at 65°C for 1 h. The tunnel’s height was 28 μM and its width at the worm’s nose space was 24 μM.

#### Dual-olfactory chip fabrication

The dual-olfactory chips were designed in AutoCAD 2022 and converted to CleWin to create the digital mask for fabrication with Mask-less Aligner (Heidelberg MLA150). First, we cleaned the silicon wafer with propanol and acetone, and then rinsed it with MQ-water, dehydrated it on a hot plate at 120°C for 10 min and let it to cool down to room temperature. Second, we spun-coated 3 ml SU-8 50 photoresist (MicroChem) on the 10-cm wafer. To create a 50-micron layer, we set the spinning rate to 2000 RPM. Third, we soft-baked it at 65°C for 6 min and then gradually increased the temperature to 95°C and left it at that temperature for 20 min. Fourth, we patterned the design with MLA and then post-baked it at 65°C for 1 min and then at 95°C for 5 min. Fifth, after letting the wafer cool down at room temperature, we developed it for 6 min and then rinsed it with propanol. Lastly, we dried the mold with nitrogen and hard-baked it at 120°C for 2 h.

#### PDMS preparation

We mixed Sylgard-184 and reagent at a ratio of 15:1 to obtain PDMS (polydimethylsiloxane, Dow Corning). Next, we degassed it with a desiccator for 1 h to create a fully transparent mixture. Before casting PDMS, we coated the surface of the mold with the gas of Chlorotrimethylsilane (puriss., ≥99.0% (GC) SIGMA-ALDRICH) inside a hood to facilitate the peeling process. We poured PDMS into a 1-cm-thick mold and then cured it in an oven at 85°C for 90 min. Next, we created inlets and outlets in the PDMS with a 0.75 mm round punch (0.035 x 0.025 x 1.5 304 SS TiN Coated, SYNEO EUROPE LTD.).

#### Chip fabrication

We activated the surfaces of a glass cover slip (#1.5) and the PDMS with O2-plasma for 30 sec at 30W and then bonded them together with a gentle touch and then left the chip on a at 120°C hot plate for 10 min to strengthen the bonding. Ultimately, we connected the tubes to the inlets and outlets of the chip with hollow stainless-steel pins.

#### Experimental set-up and operation

The microfluidic chips were operated using two pumps that control the flow of buffer and stimulus into the microfluidic chips. The solutions were pushed through PVC tubes and stainless-steel connectors into the tunnels of the chip. We determined the arrival of the stimulus to the worm with a manual switch (or two switches when the dual-olfactory chip was used). Tubes were replaced between experiments and the connectors were cleaned with ethanol. The flow rate during the experiments was ~0.005 ml/min. Loading the worm into the chip was done by placing the worm in a drop of S-basal buffer, sucking the drop with a 1 ml syringe and inserting it into the relevant inlet of the chip. To prevent movement, 10 mM Levamisole was added to all solutions. To visualize the proper delivery of a stimulus to the worm, 50 μM rhodamine B was added only to the stimulus. If the worm moved or the flow was incorrect, the file was discarded, and a second trial was performed with the same worm. No more than two trials were done with the same worm. Imaging was done with a Zeiss LSM 880 confocal microscope using a 40x magnification water objective. The imaging rate was 6.667 Hz, the total imaging duration was 1 min, and the stimulus duration was 20 sec. A stimulus was given at 20–40 sec from imaging initiation. For analysis, the GCaMP6s fluorescence intensity was measured using FIJI. All the files were exported as tiff files, ROIs of the somas were drawn manually to best represent the signal and their mean gray values were exported. The data analysis was performed using MATLAB. For each worm, the baseline fluorescent level (F0) was calculated by averaging the mean gray values of 66 frames (10 sec) before stimulus delivery. Then, for each frame, the ΔF was calculated by subtracting F0 from the value of that time point, and the result was divided by F0, to normalize the differences in the fluorescence baseline levels between individuals (ΔF/F0). The first 66 frames in each recording, i.e., those prior to the frames used for normalization, were discarded from the data. Thus, in the finalized dataset, there are 50 sec of recording and the stimulus appears as given between 10–30 seconds.

All statistical comparisons were done on the normalized data. The moving mean of each animal’s recording data was computed across 7 frames (~1 sec). Peak responses were calculated as the difference between the maximal values for excitation or minimal values for inhibition (during 20 sec of stimulus) and the starting value when the stimulus was given.

### DiD Staining

Worms were washed with M9 buffer and then incubated in 1 ml M9 and 5μl DiD dye at ~25 RPM and tilted for 1 h. The worms were then transferred to a fresh plate and let to crawl on a bacterial lawn prior to imaging. The PHA neuron was identified as the more anterior cell and the PHB neuron as the more posterior cell filled with dye.

### Behavioral repulsion assay: Tail drop

The tail drop avoidance assay was done as described previously^19,20^. All the assays were performed on 1-day adult hermaphrodites. Briefly, worms were given 10 min to habituate on a foodless NGM experiment plate, and then underwent 8–10 repellent stimulations with 2 min intervals between stimuli. A small drop of the repellent (glycerol in S basal was placed on the agar near the tail of a forward-moving animal, using a 10 μl glass-calibrated pipette (VWR International) was pulled by hand on a flame to create two needles with a reduced diameter. The pipette was mounted onto a holder with a rubber tube, operated by mouth. A day before the experiment, unseeded NGM plates were taken out of 4°C storage, dried at 37°C for 2 h, and then left on the bench. Scoring for each trial was binary (1 for reversal, 0 for no reversal). The average score of all trials is the avoidance index for each animal.

### Behavioral – optogenetics

For optogenetic activation, we used worms expressing ChR2, only in ASH (*gpa-13p∷FLPase, sra-6p∷FTF∷ChR2∷YFP* (*ljIs114*)^38^ and/or PHB (pHS9 - *gpa-6p∷Chr2∷mCherry*), with a genetic background of *lite-1(ce314)*. L4 hermaphrodites were picked a day before the experiment and separated into control and experiment groups. They were transferred to newly seeded plates with 300 μl OP50 that was concentrated 1:10. ATR (all-trans-retinal) was added only to the experimental groups’ plates, to a final concentration of 100 μM. As ATR is sensitive to light, all plates were handled in the dark. Tracking and optogenetic stimulations were done on freshly seeded NGM test plates: on the day of the experiment, the plates were seeded with 30 μl OP50. ATR was added to the experimental group’s plates. A group of up to 10 worms was transferred to the test plates. After 10 min of habituation, the worms were tracked for 69 sec with five 2-sec 470 nm LED (1.6 mW/mm^2^) activations each, and 10 sec ISI. Each recording started with 10 sec without LED. Analysis was performed manually. If the worm reversed during a 3 sec window (2 sec LED duration + one additional second), it received a score of one, otherwise a score of zero. The five results of each worm were averaged to a number between zero and one. Worms that left the FOV of the camera were excluded. Speed measurements were extracted from WormLab.

### Computational model

We adapted the model used in Pechuk et al.^28^ and simulated the response of the nociceptive circuit in the hermaphrodites to sensory stimulation. To the cells that were previously used (ASH, AVA, AVB, AVD, PVC, A, and B), we added the sensory cells in the tail, PHA and PHB. A movement’s direction was determined by the difference in the activation of the two motor neuron groups^28,66^. Connectivity data (taking into consideration the connections’ strength) was taken from Cook et al^24^.

To find possible polarity configurations of this circuit, we explored all the combinations of inhibition and excitation in its 21 chemical synapses, resulting in 2^21^ configurations. To account for possible variability in the biophysical parameters^28,67^, we tested all the polarity configurations on 50 different parameter sets (50 ⋅ 2^21^ ≈ 10^8^ combinations overall). The parameter sets were randomly sampled from sets that met all the behavioral and physiological conditions in hermaphrodites, as previously defined^28^. Briefly, the behavioral conditions included backward movement after a strong sensory stimulus and forward movement otherwise, a reasonable range of membrane potentials and time constant, and the ability of neurons to return to rest after the sensory stimulus ended (see “behavioral conditions” in^28^ for additional details). The physiological conditions included anticorrelated activity between cells promoting backward movement (A and AVA) and cells promoting forward movement (B and AVB), in agreement with the expected behavior (see “physiological conditions” in ^28^ for additional details).

We simulated the membrane potential of the neurons for 15 sec, while the sensory stimulus was delivered for 5 sec, starting from the fifth sec. Basal inputs to the interneurons were simulated throughout the simulation. We used two sensory stimuli: one delivered only to ASH, and another delivered to all 3 sensory neurons (simulated separately). Noise was not simulated in the inputs or in the synaptic strength values. In both sensory stimuli, the sets had to meet all the behavioral and physiological conditions specified above^28^. On top on those conditions, we added a demand for tail antagonism^19^, i.e., the difference in activation between the motor neurons had to be smaller after a sensory stimulus to all sensory neurons, compared to stimulation of ASH alone. This condition reflects the antagonistic effect of the tail sensory neurons on the head sensory neuron. For each synapse, we summed over the results in sets of parameters and polarity that met all the conditions, checking the proportion in which it was inhibitory vs. excitatory.

## DATA AVAILABILITY

The authors declare that all the data generated or analyzed during this study are included in this published article (and its supplementary information files). Source data are provided with this paper.

## ACKNOWLEDGEMENTS

We thank members of the Oren-Suissa lab for their critical insights regarding the manuscript. Some strains were provided by the CGC, which is funded by the NIH Office of Research Infrastructure Programs (P40 OD010440).

MOS acknowledges financial support from the European Research Council ERC-2019-STG 850784, Israel Science Foundation grant 961/21, Dr. Barry Sherman Institute for Medicinal Chemistry, Sagol Weizmann-MIT Bridge Program and the Azrieli Foundation. MOS is the incumbent of the Jenna and Julia Birnbach Family Career Development Chair. MK acknowledges financial support from the ERC (MechanoSystems, 715243), HFSP (CDA00023/2018), Spanish Ministry of Economy and Competitiveness through the Plan Nacional (PGC2018-097882-A-I00), FEDER (EQC2018-005048-P), “Severo Ochoa” program for Centres of Excellence in R&D (CEX2019-000910-S; RYC-2016-21062), Fundació Privada Cellex, Fundació Mir-Puig, and from Generalitat de Catalunya through the CERCA and Research program (2017 SGR 1012).

## AUTHOR CONTRIBUTIONS

AG, VP and SPK conducted and analyzed the experiments. GG built the computational model. AG, SK and MK designed and fabricated the dual-olfactory chip. AG and JL analyzed the puncta’s localization. MOS supervised and designed the experiments. AG and MOS wrote the paper.

