## Supplemental data for "Integration of spatially opposing cues by a single interneuron guides decision making in *C. elegans*"

**SUPPLEMNETAL INFORMATION:**

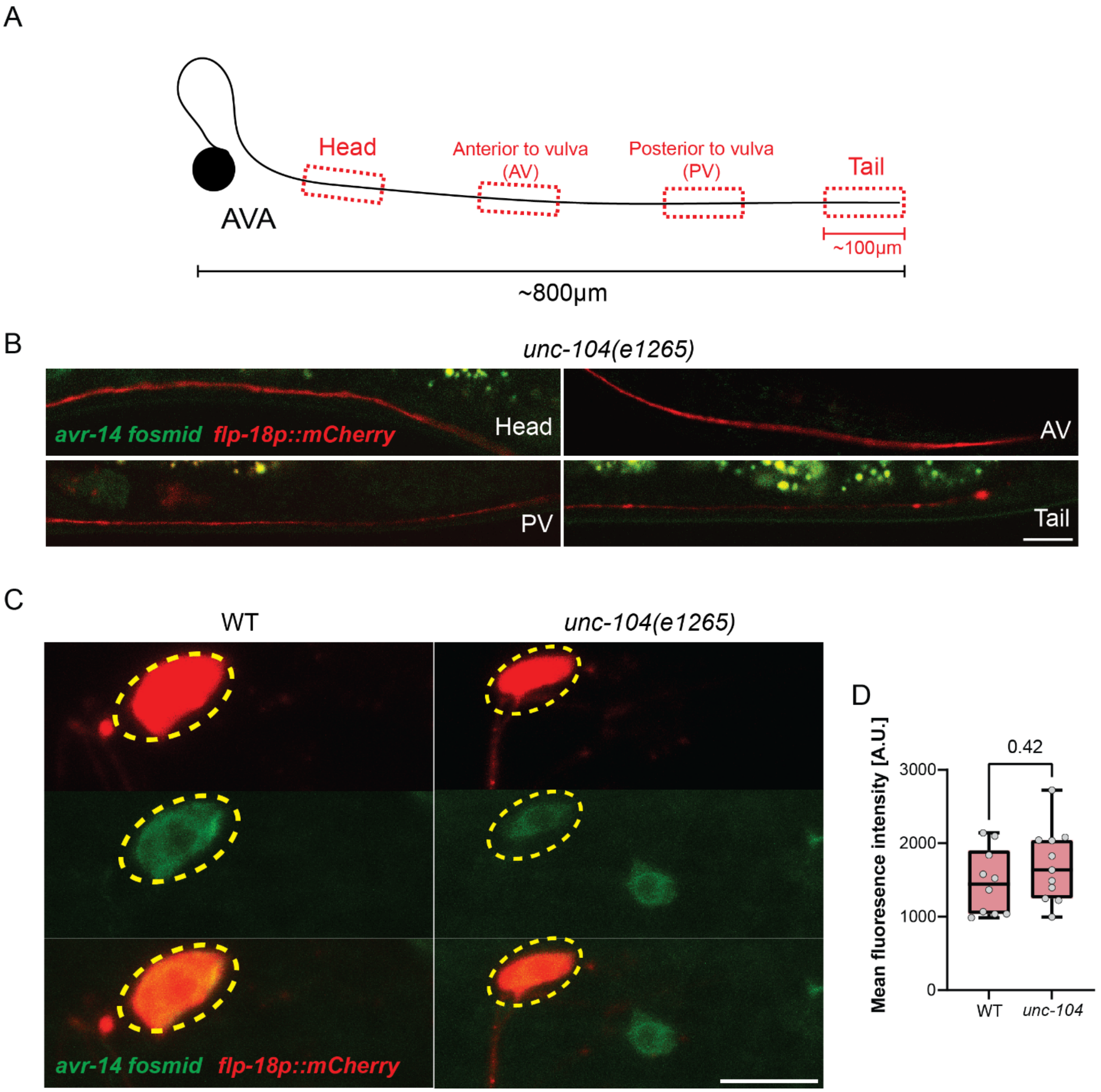

**Figure S1: The Kinesin-like protein UNC-104 is required for AVR-14 localization along the AVA process.**

(A) Schematics of the AVA process regions that were examined (red dashed-line boxes). Each examined region along the process was ~100 µm in length. AVA process length is roughly ~800µm. (B) Representative confocal images for all four regions of the AVA process expressing a red fluorescence marker in AVA (*flp-18::mCherry*) along with GFP-fused *avr-14 fosmid* reporter. (C) Representative confocal images of AVA soma in wild-type and *unc-104(e1265)* in animals expressing red fluorescence marker in AVA (*flp-18::mCherry*) along with GFP-fused reporter of *avr-14 fosmid*. (D) Quantification of (C). n=10-11. We performed a Mann-Whitney test.

**Table 1. List of Strains used in this study**

| Strain | Description | Source |
|---|---|---|
| MOS52 | lite-1(ce314); ljIs114[Pgpa-13::FLPase, Psra-6::FTF::ChR2::YFP]; him-5(e1490)V | Pechuk et al., 2022 |
| MOS275 | etyEx73[gpa-6::ChR2::mCherry (50ng/ul) + ttx-3::GFP (30ng/ul) +pBS (20ng/ul)] ; lite-1(ce314) ; ljIs114[Pgpa-13::FLPase, Psra-6::FTF::ChR2::YFP] ; him-5(e1490)V | This study |
| MOS276 | etyEx73[gpa-6::ChR2::mCherry (50ng/ul) + ttx-3::GFP (30ng/ul) +pBS (20ng/ul)] ; lite-1(ce314) ; him-5(e1490)V | This study |
| MOS88 | etyIs1[Pgpa-13::FLPase, Psra-6::FTF::GCaMP6s]; him-5(e1490)V | Pechuk et al., 2022 |
| OH15500 | otIs669; otIs672 | Oliver Hobert's lab |
| MOS134 | etyEx35[Pflp-18::GCaMP6s, ttx-3::mCherry]; him-5(e1490)V | Pechuk et al., 2022 |
| MOS427 | ceh-14(ch3) X; him-5(e1490) V; etyEx35[flp-18::GCaMP6s, ttx-3::mCherry] | This study |
| MOS492 | kyEx5104 [pNP424 (sra-6::HisCl1::SL2::mCherry), 50 ng/µL + elt-2::mCherry, 1 ng/µL]; etyEx35[Pflp-18::GCaMP6s, ttx-3::mCherry]; him-5(e1490)V | This study |
| MOS363 | avr-14(ad1302) I; etyEx35[flp-18::GCaMP6s, ttx-3::mCherry]; him-5(e1490)V | This study |
| MOS397 | etyEx35[flp-18::GCaMP6s, ttx-3::mCherry]; him-5(e1490)V;glc-4(ok212) II. | This study |
| MOS444 | avr-14(ad1302) I; him-5(e1490) V; etyEx140[flp-18p::avr-14(cDNA-first 6 exons)::avr-14(gDNA from exon 6 to end) 30ng/ul, unc-122::GFP 30ng/ul, pBS 40ng/ul]; etyEx35[Pflp-18::GCaMP6s, ttx-3::mCherry] | This study |

| | | |
|---|---|---|
| MOS499 | nmr-1(ak4) II;glr-1(n2461) III; him-5(e1490)V; etyEx35[Pflp-18::GCaMP6s, ttx-3::mCherry] | This study |
| MOS526 | etyEx181 [MVC11 15ng/ul, pRF4 50ng/ul, nmr-1 fosmid 15ng/ul, pBS 20ng/ul] | This study |
| MOS436 | etyEx138[MVC11 10ng/ul, pRF4 50ng/ul, gpa-6::mKate2 30ng/ul, avr-14 fosmid 10ng/ul] | This study |
| MOS472 | unc-104(e1265) II; him-5(e1490) V; etyEx138[MVC11 10ng/ul, pRF4 50ng/ul, gpa-6::mKate2 30ng/ul, avr-14 fosmid 10ng/ul] | This study |
| MOS523 | etyEx180 [MVC11 10ng/ul, pRF4 50ng/ul, pAG4 Pflp-18::GLR-1::GFP 40ng/ul] | This study |
| CB4088 | him-5(e1490) V | Caenorhabditis Genetics Center |
| MOS376 | nmr-1(ak4) II ; glr-1(n2461) III; him-5(e1490)V | This study |
| MOS527 | him-5(e1490) V; avr-14(ad1302) I | This study |
| MOS459 | him-5(e1490) V; avr-14(ad1302) I; nmr-1; glr-1(n2461) III | This study |
| MOS623 | avr-14(ad1302) I; him-5(e1490) V; etyEx140[flp-18p::avr-14(cDNA-first 6 exons)::avr-14(gDNA from exon 6 to end) 30ng/ul, unc-122::GFP 30ng/ul, pBS 40ng/ul]; | This study |
| MOS594 | him-5(e1490)V; lite-1(ce314)X;glr-1(n2461) III; nmr-1(ak4) II; etyEx73[gpa-6::ChR2::mCherry (50ng/ul) + ttx-3::GFP (30ng/ul) +pBS (20ng/ul)] ; ljIs114[Pgpa-13::FLPase, Psra-6::FTF::ChR2::YFP]X | This study |
| MOS562 | him-5(e1490)V; lite-1(ce314)X;glr-1(n2461) III; nmr-1(ak4) II; avr-14(ad1302) I;etyEx73[gpa-6::ChR2::mCherry (50ng/ul) + ttx-3::GFP (30ng/ul) +pBS (20ng/ul)] ; ljIs114[Pgpa-13::FLPase, Psra-6::FTF::ChR2::YFP]X | This study |
| MOS563 | him-5(e1490)V; lite-1(ce314)X;glr-1(n2461) III; nmr-1(ak4) II; avr-14(ad1302) I;etyEx73[gpa-6::ChR2::mCherry (50ng/ul) + ttx-3::GFP (30ng/ul) +pBS (20ng/ul)] ; ljIs114[Pgpa-13::FLPase, Psra-6::FTF::ChR2::YFP]X; etyEx140[flp-18p::avr-14(cDNA-first 6 | This study |

| | |
|---|---|
| | exons)::avr-14(gDNA from exon 6 to end) 30ng/ul, unc-122::GFP 30ng/ul, pBS 40ng/ul]; |
